# Sacbrood viruses and select Lake Sinai virus variants dominated *Apis mellifera* colonies symptomatic for European foulbrood

**DOI:** 10.1101/2024.03.11.584375

**Authors:** Poppy J. Hesketh-Best, Peter D. Fowler, Nkechi M. Odogwu, Meghan O. Milbrath, Declan C. Schroeder

## Abstract

European foulbrood (EFB) is a prevalent disease of the European honey bee (*Apis mellifera*) in the US, which can lead to colony decline and collapse. The bacterial components of EFB are well studied, but the diversity of viral infections within infected colonies has not been explored. Here we use meta-transcriptomics sequencing of 12 honey bee hives, symptomatic (+) and asymptomatic (-) for EFB to explore viral infection associated with the disease. We identified 41 viral genomes, belonging to two families, with a predominant occurrence of 34 genomes observed in colonies with severe EFB. This is in contrast to fewer, and a complete absence of *Dicistroviridae* genomes, recovered from healthy colonies (7 genomes). Identified viruses included multiple lineages previously reported in honey bees, namely Lake Sinai virus, Deformed wing virus, Sacbrood virus, Black queen cell virus, and Israeli acute paralysis virus.

We observed specific Lake Sinai virus clades associated exclusively with EFB+ or EFB-colonies, in addition to EFB-afflicted colonies that exhibited an increase in relative abundance of sacbrood viruses. Multivariant analyses highlighted that a combination of site and EFB disease status influenced RNA virome composition, while EFB status alone didn’t significantly impact it, presenting a challenge for comparisons between colonies kept in different yards. These findings contribute to the understanding of viral dynamics in honey bee colonies compromised by EFB and underscore the need for future investigations to consider viral composition when investigating EFB.

**Importance:** This study on the viromes of honey bee colonies affected by European foulbrood (EFB) sheds light on the dynamics of viral populations in bee colonies in the context of a prevalent bacterial brood disease. The identification of distinct Lake Sinai virus and Sacbrood virus clades associated with colonies with severe EFB suggests a potential connection between viral composition and disease status, emphasizing the need for further investigation into the role of viruses during EFB infection. The observed increase in sacbrood viruses during EFB infection implies a possible viral dysbiosis, with potential implications for honey bee brood health. These findings contribute valuable insights for apicultural practices, offering a foundation for future research aimed at understanding and mitigating the impact of bacterial and viral infection in commercial honey bee operations and the management of EFB.

## Introduction

European honey bees (*Apis mellifera*) are hosts to a wide diversity of pathogenic viruses and bacteria which can contribute to colony illness and death. The honey bee industry is a crucial contributor to global agricultural systems, and in the US alone, honey bees are estimated by the Department of Agriculture to provide an annual value of 18 billion USD (1). However, parasites and diseases have significantly impacted bee health in recent years. Data from 2022 provided by the Bee Informed Partnership shows that US commercial beekeepers experienced 46.1% loss in their colonies from 2019 to 2020 (2). Many factors can contribute to colony loss, such as adult bee physiology, pathogen loads, and pesticide levels (3, 4). European foulbrood, commonly referred to as EFB, is a highly prevalent bacterial disease of honey bee larvae within the US, with *Melissococcus plutonius* identified as the primary pathogen based on genomic studies, field observations, and laboratory testing (5, 6). Two studies have recently focused on EFB in the US and both have noted high rates of disease and risk to colony growth particularly after blueberry field pollination (7, 8). Honey bees infected with EFB can exhibit high mortality, but disease monitoring is complicated by *M. plutonius* being present in both symptomatic and asymptomatic colonies (9, 10). Currently, the only approved treatment in the US for EFB is the antibiotic Oxytetracycline, for which antimicrobial-resistant strains have been detected in North American honey bee colonies (11). Not only are better treatments urgently needed, but improved fundamental knowledge of the disease and its progression, in addition to the onset of secondary infections is also necessary to manage this disease.

The stresses associated with EFB may leave colonies susceptible to secondary infections. The monitoring and understanding of the range of secondary pathogens that can complicate disease management and treatments can be as important as diagnosing the primary infection. EFB secondary bacterial infective agents include *Enterococcus faecalis* and *Paenibacillus alvei*, and are well described (6, 7, 9). Secondary viral infections have been largely overlooked, and comparisons between EFB symptomatic (+) and asymptomatic (-) colonies are unexplored. We previously found no correlation or association between the viral loads of the species *Iflavirus aladeformis*, also known by the virus name Deformed wing virus (DWV), and EFB. This observation is not surprising due to the efficient control of the ectoparasitic mites, *Varroa* spp., that vectors DWV (12–14). DWV symptoms typically include wing deformities, and decreased body size in infected larvae (15), while more commonly subclinical infections have been associated with impaired cognitive function and foraging performance (16–18). *Triatovirus nigereginacellulae* or Black queen cell virus (BQCV), causes larval death and decay in queen cells, but infection may be inapparent in worker brood (19, 20). *Iflavirus sacbroodi*, or Sacbrood virus (SBV), is often associated with brood that fails to pupate and instead develops in a pre-pupal-shaped sac with a shrunken head and may be inapparent in infected worker bees (21, 22). The simultaneous occurrence of brood-related bacterial and viral diseases may exert confounding effects on the overall health of the hive. This complexity is particularly evident in diagnostic processes, as exemplified by the case of *Apis cerana* bees in India, where the symptoms of Thai Sacbrood virus (TSBV) were mistaken for EFB symptoms (23). The symptoms of respiratory viral infections in humans can leave the afflicted individual compromised to secondary bacterial infection, which is associated with severe disease and increased mortality, this is well recorded with diseases such as pneumonia and influenza (24–26). Therefore, gaining a deeper understanding of the alterations within the honey bee virome resulting from EFB infection is crucial. This may help elucidate the causal relationship between severe EFB infections and composition changes to the RNA virome, as well as determine whether polymicrobial infections play a role in exacerbating disease severity.

The relationship between DWV and EFB has been briefly explored in a 2023 study that focused on commercial bees contracted for blueberry pollination, with the authors finding no relationship between DWV-A or DWV-B variants load and the severity of EFB disease (8). Given that only DWV-A and B variants were monitored for possible association with EFB, our study describes the relationships between EFB and colony based on the whole RNA virome composition. Here we used a metagenomic cDNA sequencing approach, as previously described (27, 28), on a subset of samples from 12 colonies (N=6 EFB+, and N=6 EFB-) taken from two separate yards (yard A, and yard B) that were collected as part of our previous study (8). This allowed us to explore the viral diversity of the honey bee viromes in EFB symptomatic and healthy colonies, and the relationship between EFB disease status and RNA virome composition.

## Results and Discussion

### The majority of viral genomes recovered from EFB+ colonies

We generated 6x10^8^ nanopore sequences across the 12 colonies, assembled into 300 contigs. A total of 41 fragmented and draft viral genomes were binned based on sequence composition and contig coverage with Anvi’o (29, 30). Most viral genomes were found in colonies with European foulbrood (EFB+, 34 genomes), while fewer were identified in EFB-colonies (7 genomes). Despite a comparable quantity of reads generated between colonies from both yards and EFB+/EBF-colonies, yard B colonies that were EFB-still had an overall smaller proportion of viral reads taxonomically classified by Kaiju (Table 1, Supp. Data File 1). Viral reads overall contributed 0.5-4.8% of the total cDNA sequenced, while *Melisoccocus plutonius,* the primary candidate as the causative agent of EFB, contributed between 0.001 and 0.028% of the dataset. Notably, *M. plutonius* transcripts were absent, or very low (∼0.001%), in EFB-colonies. Between 8-18% of the reads were classified as Eukaryotic and the majority as unclassified, 70-87%. This is likely a result of Kaiju being specialized for microbial classification (31), and we confirmed this by mapping the reads to the *A. mellifera* genome (GCF_003254395.2, accessed Feb 2022), which estimated that the majority of the reads (70-89%) belong to the host (Table 1; Supp. Data File 1).

**Table 1.** Summary data generation for Oxford Nanopore Technologies sequencing reads by taxonomic classification and viral genome assembly. Shown here are only reads that met the minimum quality threshold, 200 bp and Q >9. (EFB, European foulbrood; †, Apis classified reads were estimated by mapping, Minimap 2, to the *A. mellifera* genome due to low success in eukaryotic classification with Kaiju).

| EFB | Yard | Foraging | Average reads per sample | Average *A. mellifera*† (%) | Average prokaryotic reads [*M. plutonius*] (%) | Average Unassigned | Average viral reads (%) | Total Viral genomes |
| --- | --- | --- | --- | --- | --- | --- | --- | --- |
| - | Yard A | Blueberry yard | 199,596 | 81.735 | 4.576 [0.000] | 10.291 | 3.398 | 6 |
|  | Yard B | Holding yard | 198,683 | 87.680 | 3.581 [0.000] | 8.158 | 0.581 | 1 |
| + | Yard A | Blueberry fields | 168,276 | 74.588 | 6.858 [0.014] | 13.727 | 4.827 | 19 |
|  | Yard B | Holding yard | 225,433 | 83.915 | 4.716 [0.006] | 8.720 | 2.649 | 15 |

Viral contigs were taxonomically identified using Kaiju against the Reference Viral Database (RVDB) (31, 32), and phylogenetically confirmed for distinguishing variants, particularly DWV-A and DWV-B (Fig. 1-3, and Supp. Fig. 1-2). Viral genomes were taxonomically identified as follows: 13 *Sinaivirus* spp. (Lake Sinai virus/LSV; Fig. 1), 11 *I. aladeformis* (DWV; Fig. 2), seven *I. sacbroodi* (SBV, Fig. 3), five *T. nigereginacellulae* (BQCV, supp. Fig. 1), and five *Aparavirus israelense* (Israeli acute paralysis virus/IAPV; Supp. Fig. 2). Five of these genomes were regarded as high-quality by an assessment performed with CheckV (33), 28 genomes were medium-quality and seven low-quality genomic fragments (Supp. Data File 2). None of the genomes were estimated to have any contamination, and low-quality genomes were limited due to their short lengths (<50% estimated completeness).

**Fig 1.**
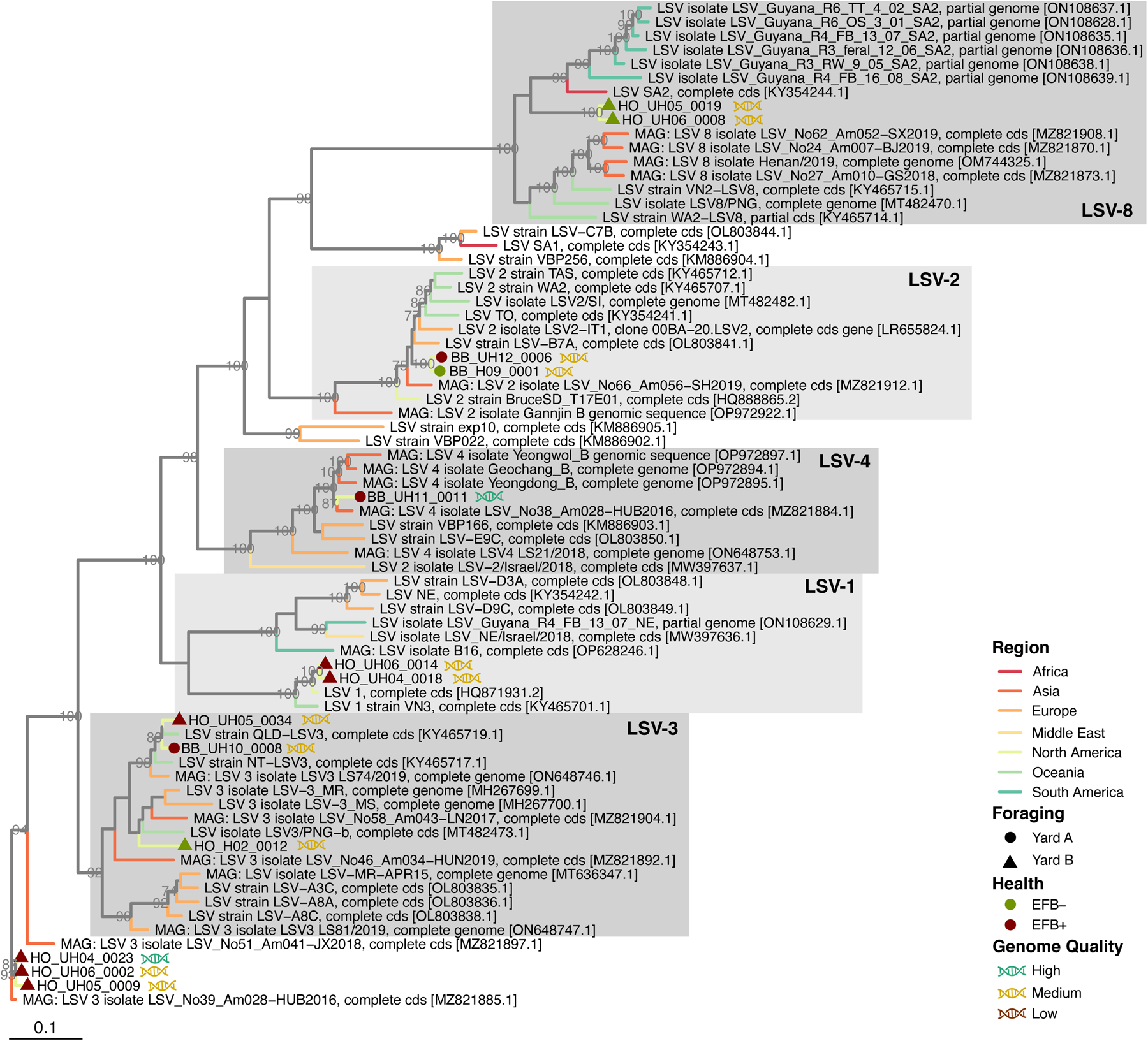
Whole-genome maximum-likelihood phylogeny of viral genomes from EFB symptomatic and asymptomatic bee viromes for Lake Sinai virus. Branches are coloured by the origin of viral genomes (see Supplementary Data file X for full metadata of genome used in phylogenies). Branch support values are from left to right, bootstraps from 1,000 replicates are reported as a proportion out of 100. Viral genomes accessed from NCBI are denoted by their accession numbers, while genomes assembled in this study are annotated with their EFB health record, their holding yard, and genomes by their quality.

**Fig 2.**
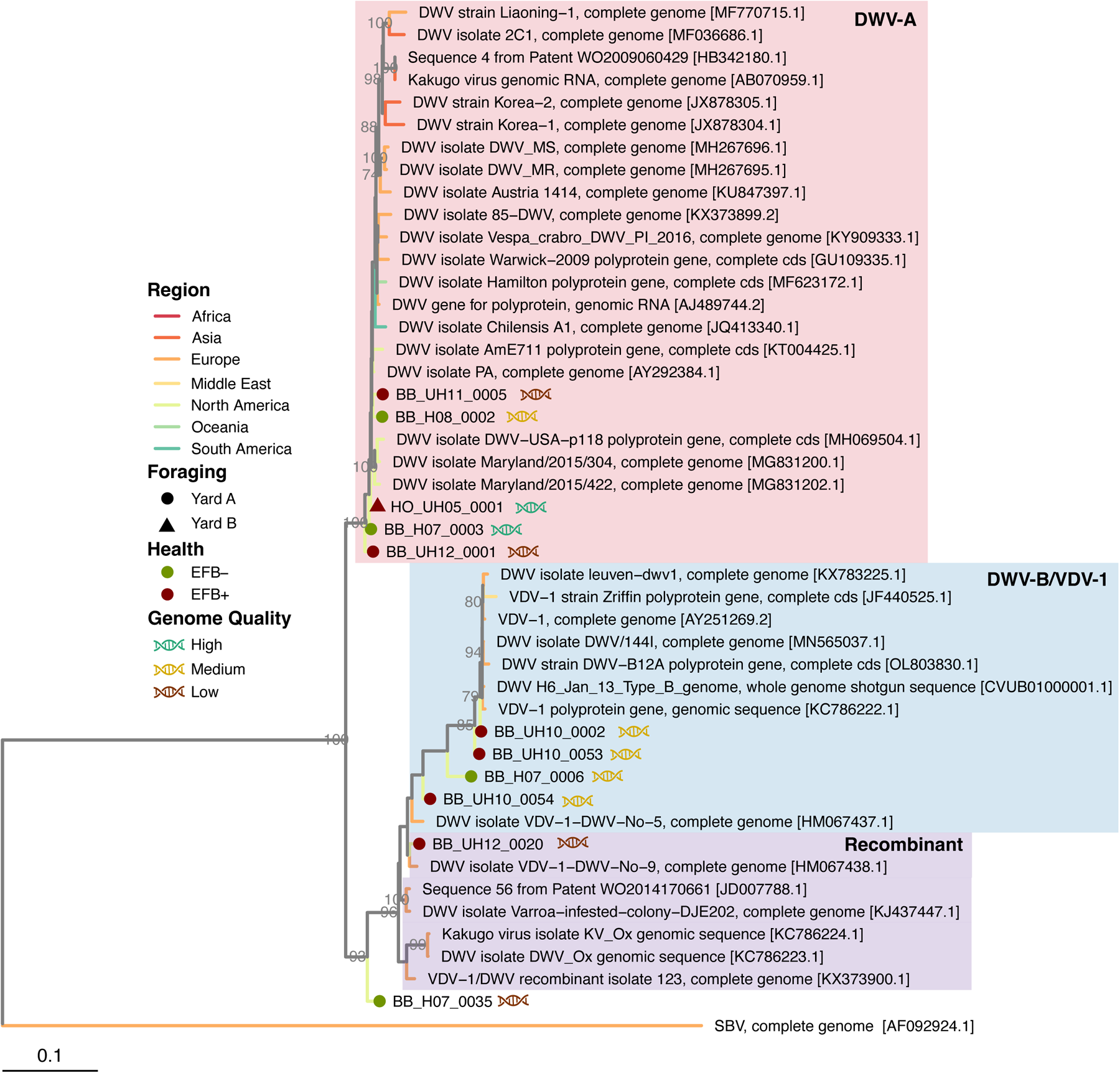
Whole-genome maximum-likelihood phylogeny of viral genomes from EFB symptomatic and asymptomatic bee viromes for deformed wing virus. Branches are coloured by the origin of viral genomes (see Supplementary Data file X for full metadata of genome used in phylogenies). Branch support values are from left to right, bootstrap from 1000 replicates reported as a proportion out of 100. Viral genomes accessed from NCBI are denoted by their accession numbers, while genomes assembled in this study are annotated with their EFB health record, their holding yard, and genomes by their quality.

**Fig 3.**
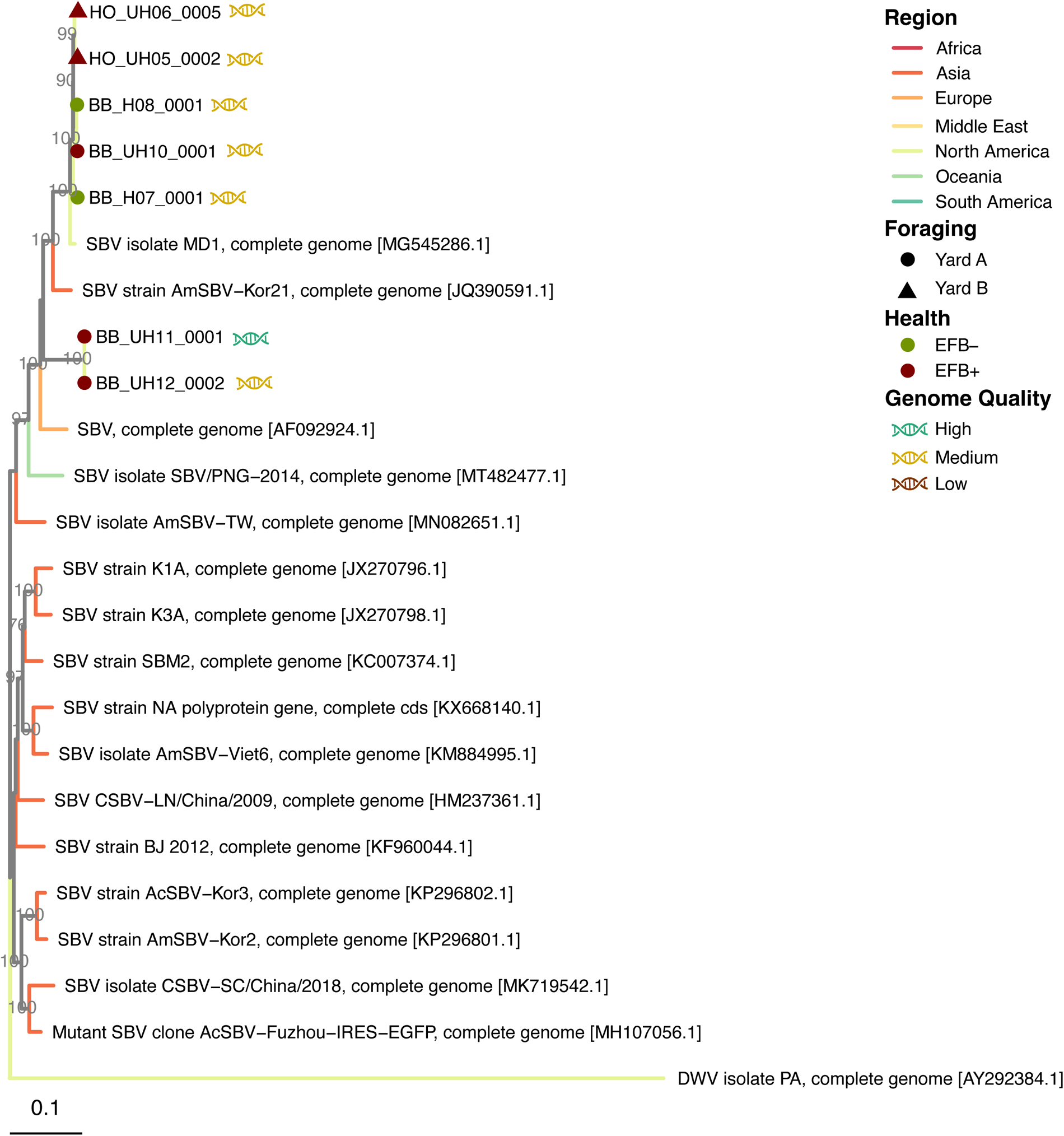
Whole-genome maximum-likelihood phylogeny of viral genomes from EFB symptomatic and asymptomatic bee viromes for sacbrood virus. Branches are coloured by the origin of viral genomes (see Supplementary Data file X for full metadata of genome used in phylogenies). Branch support values are from left to right, bootstrap from 1000 replicates reported as a proportion out of 100. Viral genomes accessed from NCBI are denoted by their accession numbers, while genomes assembled in this study are annotated with their EFB health record, their holding yard, and genomes by their quality.

### Lake Sinai virus and Sacbrood virus load and diversity distinguish EFB+ and EFB-colonies

Genomes representing LSV were the most abundant (13 genomes, Fig 4A) and represented five described clades of LSV: two LSV-1 genomes, two LSV-2, three LSV-3, one LSV-4, two LSV-8, and three genomes that did not cluster with any of the presently described LSV clades (Fig. 1). Notably, genomes of LSV-1, LSV-4, and LSV-3 were exclusively present in EFB+ colonies, whereas LSV-2 genomes were detected in both EFB+ and EFB-colonies. LSV-8 genomes were only recovered from yard B colonies that were EFB-, and finally LSV-2 and LSV-4 genomes from yard A colonies. This is additionally supported by rank-abundances of viral genomes between the sites, where we observed low-ranked LSV-8 genomes in the yard A colonies and highly ranked in yard B colonies (Fig. 4B). Furthermore, the two LSV-8 variants clustered with a clade within LSV-8 that originated from South Africa and Africanized honey bees in Guayana (27). The origin of Africanized bee in South America was South Africa, which indicates that these colonies, which originated from Florida, may have been exposed to Africanized bee, which has previously been reported in Southern US states (34, 35).

**Fig 4.**
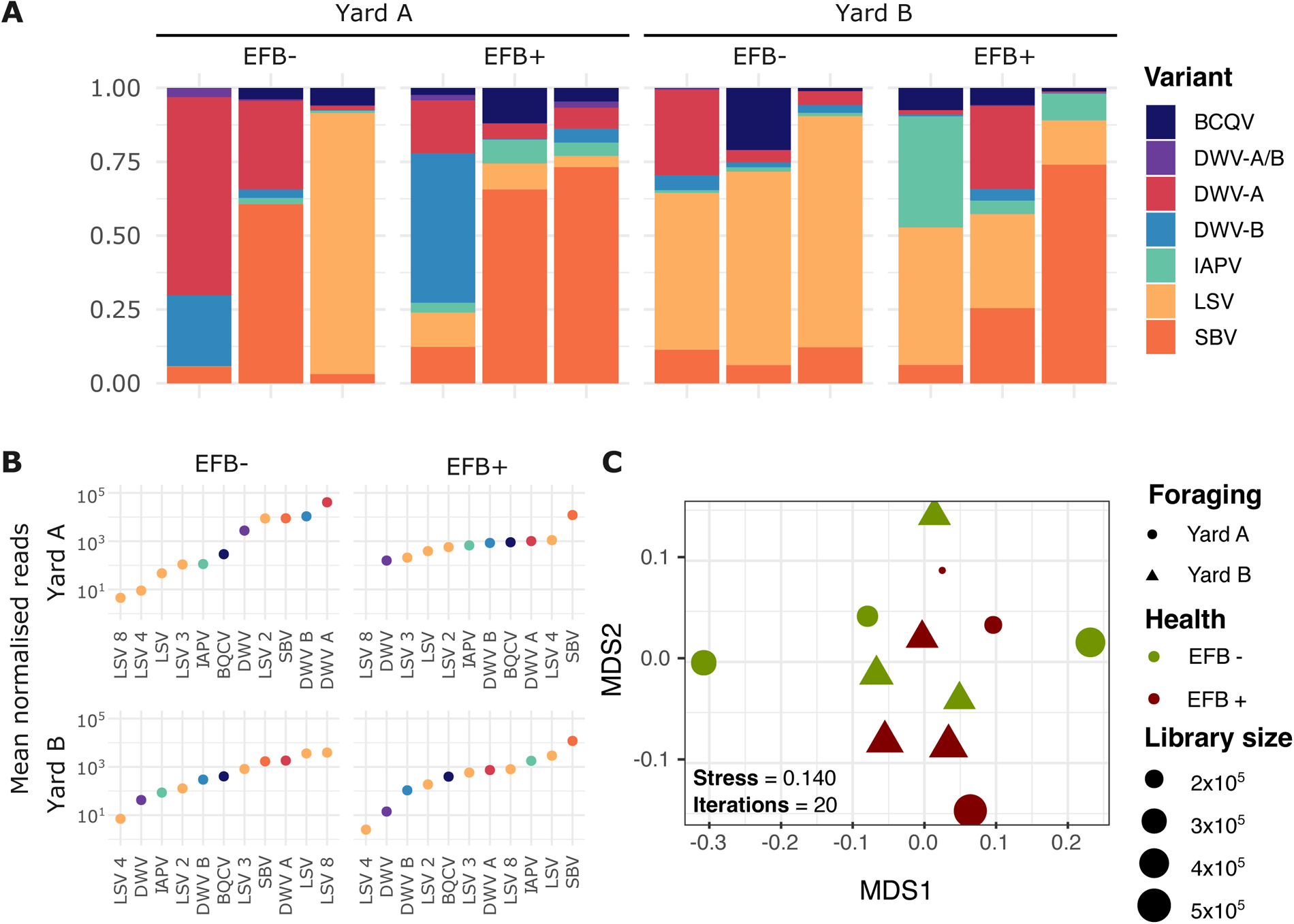
(A) RNA virome community composition nested by EFB symptoms and colony sites. (B) Rank abundance (mean) of each viral genome (grouped by lineage and variant). (C) MDMS plots of a Bray-Curtis dissimilarity matrix of cumulative sum scaled normalized read counts to the viral genome collection, colors represent the EFB status, shape colony yards and datapoint size represent the size of the sequencing library.

There remains a very limited understanding of the impact LSV has on colony health. Still, they are reported as globally widespread (27, 36). LSV-2 in particular has been implicated in colony health for migratory bee colonies (37). Overall LSV were the most abundance genomes in EFB-colonies, as measured by read-recruitment (Fig. 4). What impact LSV has on EFB disease has not been investigated, but the observations here indicate a phylogenetic distinction between EFB+ and EFB-infecting LSVs genomes (Fig 1).

In this dataset, we observed during EFB+ infection an increase in the fraction of virome reads assigned to DWV and IAPV genomes in EFB+ colonies, while BQCV does not differ between EBF+ and EFB-colonies. Furthermore, our study noted a rise in reads recruitment to binned SBV genomes in EFB+ colonies. In prior research, where SBV has not consistently correlated with colony health decline (38), our findings indicate a prevalence of SBV in EFB+ colonies (Fig 4B). The results in this study are impeded by fewer viruses recovered from EFB-colonies, as compared to EFB+ colonies, and should be taken cautiously lacking any quantitative data on viral density for all the viral lineages reconstructed from the RNA viromes. However, we do observe a distinct clade of SBVs, comprised only of genomes recovered from EFB+ colonies, while genomes from EFB-colonies, along with additional EFB-, clustered closely together with North American variants (Fig. 3). Our observation might indicate a form of viral dysbiosis in the event of EFB infection that shifts the community towards one abundant in SBV. This is particularly interesting as EFB is a brood disease like SBV, and exploring the impact on brood viral composition during EFB may reveal more. While SBV can be asymptomatic in adult bees (39–42), high densities in hives recovering from EFB may be detrimental to the brood and the recovery of the hide during treatment for EFB.

### DWV-A and DWV-B are present in EFB+ and EFB-colonies

The data presented here lends support to the original study (Supp. Fig 3), which within the larger dataset found no difference in means of the viral loads of DWV between diseased and healthy colonies, as measured by RT-qPCR (8). Here we can add the additional support of genomic similarity between DWV genomes recovered from both diseased and healthy colonies. Of the DWV genomes assembled, five represent variant A (DWV-A) and four variant B (DWV-B) (Fig. 2). Only DWV-A genomes were classified as high-quality genomes. Two genomes of low quality clustered with DWV recombinant genomes, and due to short contig length (3.00 and 3.21 Kbp, Supp. Data File 2), taxonomy could not be fully ascertained. DWV-A genomes clustered closely with a DWV genome isolated from North American bees (Fig. 2). All DWV-B genomes from the dataset clustered closely in the tree and were recovered from yard A colonies, with all but a single genome from a colony with severe EFB symptoms, and the closest neighbors are genomes isolated from European colonies. Notably, in EFB-colonies from yard A, we observed a notable abundance of both DWV-A and DWV-B, a trend not mirrored in colonies from yard B (see Fig. 4B). In EFB-compromised colonies in yard A, there was a decrease in the ranking of DWV-A and DWV-B, with SBV emerging as the predominant genome. Despite DWV-A and DWV-B not being the dominant genomes in yard B colonies, a similar shift in dominance was evident with the onset of EFB.

### Dicistroviridae viruses have limited presence in EFB-colonies

Genomes for members of the viral family *Dicistroviridae*, which include IAPV and BQCV, were recovered only from EFB+ colonies (Supp Fig 1B). The recovered BQCV genomes were predominantly of medium or low quality, originating from EFB+ colonies and showing clustering with a nearby North American BQCV genome (Supp. Fig. 1). BQCV was found at relatively low abundances across all samples (Fig. 4A), except for a singular EFB+ colony in yard A and an EFB-colony in yard B. However, no complete genomes were obtained from the EFB-colony. Notably, BQCV genomes were exclusively recoverable from EFB+ colonies, highlighting its association with this brood disease in honey bees. Prior marker gene surveys have shown BQCV as prominent in queen bees (43), and have been associated with colony losses in multiple countries for both *A. cerana* and *A. mellifera* (44, 45), and globally widespread in bee operations (46–48). Unfortunately, there is little prior work exploring the interactions of brood-afflicting viruses and EFB.

IAPV is a member of the AKI virus complex assembled from the dataset (49), and ranked mid to low abundance in all situations except for EFB+ yard B colonies (Fig. 4B), where it ranked highly just below SBV and LSV. A previous marker gene study has shown that bees exposed to migratory management during adulthood had increased levels of the AKI virus complex (50). Although the colonies analyzed in this study were relocated from Florida to Michigan for blueberry pollination, the absence of information about the colony’s virome composition before transportation limits our ability to assess the potential impacts of the relocation. The absence of knowledge regarding the colony virome composition before transportation for this dataset complicates our understanding of the long-term changes the colonies have undergone as a result of transportation and the onset of EFB symptoms. Future work addressing the relationship between viral infections and EFB may need to consider such variables.

### Viral composition varies with colony yards

Despite the constraints posed by the small sample size (n=12), insights into community composition were gained through an analysis of variance using distance matrices (ADONIS) (Fig. 4C). The results of the ADONIS test indicate that the EFB health status (EFB+ or EFB-), yard (A or B), and virus species, as well as their interactions, collectively explain a proportion of the variation observed in the dataset (supp. table 1). Specifically, virus species demonstrate a statistically significant effect (F = 3.709, *p* = 0.001). Additionally, the interaction between EFB health status and virus species (F = 3.171, *p* = 0.001), and yard and virus species (F = 2.342, *p* = 0.007) also exhibit significant effects. However, the main effects of EFB status and yard individually show only marginal significance (EFB: F = 2.438, *p* = 0.058; Yard: F = 1.127, *p* = 0.302), suggesting that while they may contribute to the model, their impact is not as pronounced as the virus species and its interactions. Overall, the results underscore the importance of considering the combined effects of EFB, Yard, and virus species, as well as their interactions, in explaining the observed patterns in the dataset. These findings, while informative, must be interpreted cautiously due to the inherent limitations of the small sample size and the observed variation between colonies kept in different yards. Furthermore, distinct management approaches to colonies between the two yards may additionally contribute to the variation. The samples utilized in this study were collected during advanced stages of the disease, limiting our understanding of the temporal changes to the virome composition during disease progression. Colonies within the same yard, acting as replicas, exhibit distinct viral compositions. This may influence the study design, and underscores the importance of considering this factor, especially when comparing bee colonies held in multiple locations. Understanding whether viral infections worsen EFB severity underscores the importance of early detection and continuous monitoring of the viral consortium throughout.

## Conclusion

Here we report the RNA viromes from honey bee colonies at two sites from a commercial operation in Michigan. In this study, we observe a distinct community structure in the viromes of colonies kept in two distinct yards, regardless of EFB disease. The original study, from which this dataset was subsampled, focused only on DWV and detected no relationship between hygienic behavior and the progression of EFB. We observed that SBV were dominant in EFB+ colonies, while LSV were highly prevalent in EFB-and EFB+ colonies, with specific clades associated with EFB+ and EFB-colonies. We further observe large variations between colonies from two different yards. This study is limited in scope and size and requires expansion, but it highlights the need for a better understanding of the effects of bacterial brood diseases on the viral dynamics in infected colonies.

## Methods

### Experimental design

The full study design, acquisition and storage of honey bees can be found in the original publication (8), in summary, 60 hives from a large commercial beekeeping operation in Southern Michigan were split into two groups. Yard A consisted of colonies (n=6) used for blueberry pollination and yard B colonies (n=6) were kept in a large holding yard away from crop pollination but in the same region of Michigan (Supp. Data File 1). Colonies were monitored between May and June for European foulbrood. Adult nurse bees (approximately 150 individuals) were collected from each colony at each timepoint using a sterile 60-ml centrifuge tube stored at 4°C in the field and transferred to −20 °C until processing. A subset of 12 colonies representing the two groups, of colonies showing severe symptoms of EFB and asymptomatic colonies. Pooled honey bees (n=30) were transported to the University of Minnesota on dry ice, before being stored at -80°C. European foulbrood disease was confirmed through the culture of Melissococcus plutonius from diseased colonies as previously described (8).

### RNA extraction, cDNA synthesis and nanopore sequencing

As previously described (27, 28), pooled honey bees (n=30) were dissociated with 15 mL of molecular grade water using gentleMACS™ M Tubes (Miltenyi Biotec, USA) and a gentleMACS™ Octo Dissociator (Miltenyi Biotec, USA). Bee homogenate was allowed to settle and 1 mL transferred to a centrifuged tube and centrifuged at 20,000 x g for 10 minutes at 4°C. A 200 ul aliquot was used for RNA extraction using a NucleoMag Virus kit (Macherey-Nagel, Düren, Germany) and a Magnetic Particle Processor (MPP) (KingFisher Flex, Thermo Fisher Scientific, USA). Complementary DNA (cDNA) was synthesized. For the first strand synthesis, 10 ul of viral RNA, 1 ul of N6 Primer II A (24 uM, TakaraBio, USA), 1 ul of SMARTer IIA Oligo (10 uM, TakaraBio, USA), 1 ul of 10x Template Switching Reverse Transcriptase (New England Biolabs, MA, USA) for the synthesis of the 1st strand. A 1 ul volume of Primer IIA (12 uM, TakaraBio, USA) and 1 ul of PrimeSTAR GXL polymerase (TakaraBio, USA) for the synthesis of the 2nd strand were used following manufacturer user manual instructions (TakaraBio, USA; New England Biolabs, MA, USA). The synthesized dscDNA was purified using SPRI AMPure beads according to manufacturer’s dscDNA was purified using SPRI AMPure beads. cDNA concentrations were quantified by Qubit 4 Fluorometer (QubitTM4 Fluorometer, Thermo Fisher Scientific, USA) using the dsDNA HS assay kit (Thermo Fisher Scientific, USA). Nanopore sequencing library was prepared with the Rapid Barcoding sequencing kit (SQK-RBK004) and sequenced on a FLO-MIN106 R9 flow cell for 24h. Basecalling was performed using the high-accuracy base-calling model with a minimum Q score of 9.

### Metagenomic analysis

Basecalling and demultiplexing were performed using Guppy v6.4 with the high-accuracy model. Guppy is only available to NanoPore customers through their community site (https://community.nanoporetech.com). PoreChop v0.2.4 was used to remove the nanopore barcode adapter sequences (51). Reads were error corrected and assembled using Canu v2.2 (52), using the following assembly parameters: -*nanopore maxInputCoverage=2000 corOutCoverage=all corMinCoverage=10 corMhapSensitivity=high minoverlap=50 minread=200 genomesize=5000*.

Contigs with a minimum length of 2 kbp were manually binned with the anvi’o v7.1 interactive interface (29, 30). Briefly, anvi’o profiled the contigs using Prodigal v2.6.3 (53), with default parameters, and then reads were mapped to the contig database using Minimap2 v2.24 (54). Read recruitment was stored as a BAM file using Samtools v1.19.2 (55). Anvi’o profiles each BAM file, estimating the read coverage and detection statistics of each contig. Coverage was combined into a merged profile database. The contigs database was populated with additional data, including HMMER search results against Virus Orthologous Groups (VOGs; https://vogdb.org/) in addition to the standard anvi’o HMMR profiles, NCBI COGs and KEGG KOfam database (56–59). Contig taxonomy was predicted by running Kaiju v2.9 using the non-redundant NCBI database and RVDB database (accessed 2023) on the gene calls (31, 32). Finally, the anvi’o profile was visualized for manual binning. Viral binning was guided by sequence composition similarity (visualized as a dendrogram in the Anvi’o interface), and the presence of viral HMM hits to the VOGs.

### Validation of draft and fragmented RNA virus genomes from binned contigs

Publicaly available honey bee virus genomes were downloaded from NCBI and deduplicated at 95% average nucleotide identity over 80% contig length using CD-HIT-EST (60, 61), retaining the longest genome. Publicly available genomes and genomes generated here were aligned with MAFFT v7.520 (62, 63), and IQ-TREE v2.0.3 (64, 65) was used for inference of maximum likelihood trees with 1,000 bootstrapping. Trees were visualized and metadata was added using the R package *ggtree* v3.18 (66).

### Statistical analysis and data visualization

All data was visualized using R v4.3.0, largely using *ggplot2* v3.4.4. Viral genomes were clustered based on 98% nucleotide identity, retaining the longest contig using CD-HIT v4.8.1 (Fu et al., 2012), generating a non-redundant pool of contigs that represents the viral “population” (61). Metagenomic reads were mapped to the non-redundant pool of viral genomes using Minimap2, and Samtools was utilized to estimate the number of reads mapping to each contig. An abundance matrix was calculated using the R package *MetagenomeSeq* v1.43.0 using cumulative sum scaling, a normalization method akin to median-like quantile normalization, to address disparities in sampling depth (67). For multivariate analysis of viral composition, normalized mapped read counts were utilized in conjunction with the vegan package and its metaMDS function. Additionally, an ADONIS analysis, evaluating the impact of yard and EFB symptom status, was performed using the *vegan* v2.6-7 package (68).

## Data availability

Sequence data are submitted under the BioProject ID PRJNA1072075. Nanopore sequences were submitted to the Sequence Reads Archive, under the following accession numbers: SRR27834932-SRR27834943 (Supplemental Data File 1). Scripts utilized for the assembly and binning of viral genomes and post-binning quality checks can be found in the GitHub repository: https://github.com/dmckeow/bioinf. Data tables, alignment files, tree files, viral metagenome-assembled genomes, and scripts to recreate figures and analysis are provided in the GitHub repository: https://github.com/pjhesbest/EFB_and_Foraging_honeybee_colonies-RNA_viromes_2024.

## Funding

The collection of honey bees, colony experiment work and sequencing chemistry were funded through the National Honey Board and Project Apis m. (Award 00454727). Sequencing resources were funded by D.C. Schroeder’s start-up grant from the University of Minnesota, USA. The funders had no role in study design, data collection and interpretation, or the decision to submit the work for publication.

## Author contributions

P.J. Hesketh-Best and D.C. Schroeder conceived the study. P. Fowler and M. Milbrath collected and archived bee samples used for this study. P. D. Fowler and Nkechi M. Odogwu performed the RNA extraction, cDNA synthesis and nanopore sequencing. P.J. Hesketh-Best performed the bioinformatic analysis, data curation and visualization. P.J. Hesketh-Best wrote the initial manuscript, and all authors edited and approved the final manuscript. Funding was awarded to D.C. Schroeder and M.O.G. Milbrath.

